# Chemical mechanism of allosteric and asymmetric dark reversion in a bacterial phytochrome uncovered by cryo-EM

**DOI:** 10.1101/2025.08.21.671592

**Authors:** Szabolcs Bódizs, Anna-Lena M. Fischer, Miklós Cervenak, Sayan Prodhan, Michal Maj, Sebastian Westenhoff

## Abstract

Phytochromes are light-sensitive proteins found in plants, fungi, and bacteria. They exist in two functional states, Pr and Pfr, distinguished by Z/E isomers of their bilin chromophore. The chromophore can photoswitch between these states, but also thermally converts in darkness. Despite the importance of the latter reaction, it remains unclear how it is controlled by the phytochrome. Here, we present single-particle cryo-EM measurements on the *Pseudomonas aeruginosa* bacteriophytochrome (*Pa*BphP) carried out at multiple time points during dark reversion from Pr to Pfr. These experiments resolve the structure of a PrPfr hybrid state. Surprisingly, we find that only protomer B converts back to Pfr in the hybrid, while protomer A remains in Pr. We identify structural asymmetries in the precursor Pr state, which extend from the homodimer interface to a conserved histidine (H277). The hydrogen-bonding network around the chromophore is modulated, explaining how the phytochrome gains control over the activation energy of the isomerization reaction. These findings establish that dark reversion is governed by conformational selection between two substates, whereby one is “dark-reversion ready” and the other one blocks the reaction. Moreover, we explain how the equilibrium of the states is allosterically controlled across the dimer. Together, these findings provide a structural framework for tuning phytochrome signaling lifetimes in optogenetic applications.

**Significance statement:** The dark reversion reaction of phytochromes is crucial to their signalling role in plants, bacteria, and fungi, but it is vastly understudied in terms of its chemical mechanism. It remains unsolved how the reaction can proceed at all, given that the activation energy is prohibitively high for the isomerization to occur in solution. Using time-resolved cryo-EM, we present a chemical and structural framework for understanding how the protein binding pocket regulates the reaction. Our results establish that conformational selection between two substates controls dark reversion, providing a rare example of strongly asymmetric reactivity across a dimeric protein. This opens the way for rational engineering of the lifetimes of the signaling states in phytochromes.

## Introduction

Phytochromes are universal and adaptable light sensors that regulate diverse biological processes across plants, bacteria, and fungi.^1–3^ These versatile photoreceptor proteins enable organisms to perceive and respond to light, temperature, and other environmental cues, influencing a wide array of physiological functions.^4–6^ In plants, phytochromes are central to developmental and stress response pathways, contributing significantly to fitness in natural environments and productivity in agriculture.^7–9^ Their mechanisms of action are diverse, involving changes in subcellular localization, protein-protein interactions, and crosstalk with other photoreceptors.^9–12^ In bacteria, they frequently act as histidine kinases in two-component signalling systems, although a variety of alternative output domains have evolved.^4,13^

Central to phytochrome function is their ability to reversibly switch between a red light-absorbing Pr state and a far-red light-absorbing Pfr state. This allows organisms to sense the ratio of red to far-red light in their environment. The photoswitching is initiated by light-driven Z/E isomerization of a covalently bound bilin chromophore within a conserved photosensory core module (PAS-GAF-PHY, see **Fig. 1b**).^14^ Structural studies have revealed key conformational changes between Pr and Pfr, including isomerization of the chromophore, refolding of the “PHY-tongue” motif, and rearrangements of the quaternary structure.^15–20^ More recently, single-particle cryo-electron microscopy (cryo-EM) structures have started to provide insights into full-length phytochrome architecture in near-native conditions.^21–27^

**Fig. 1.**
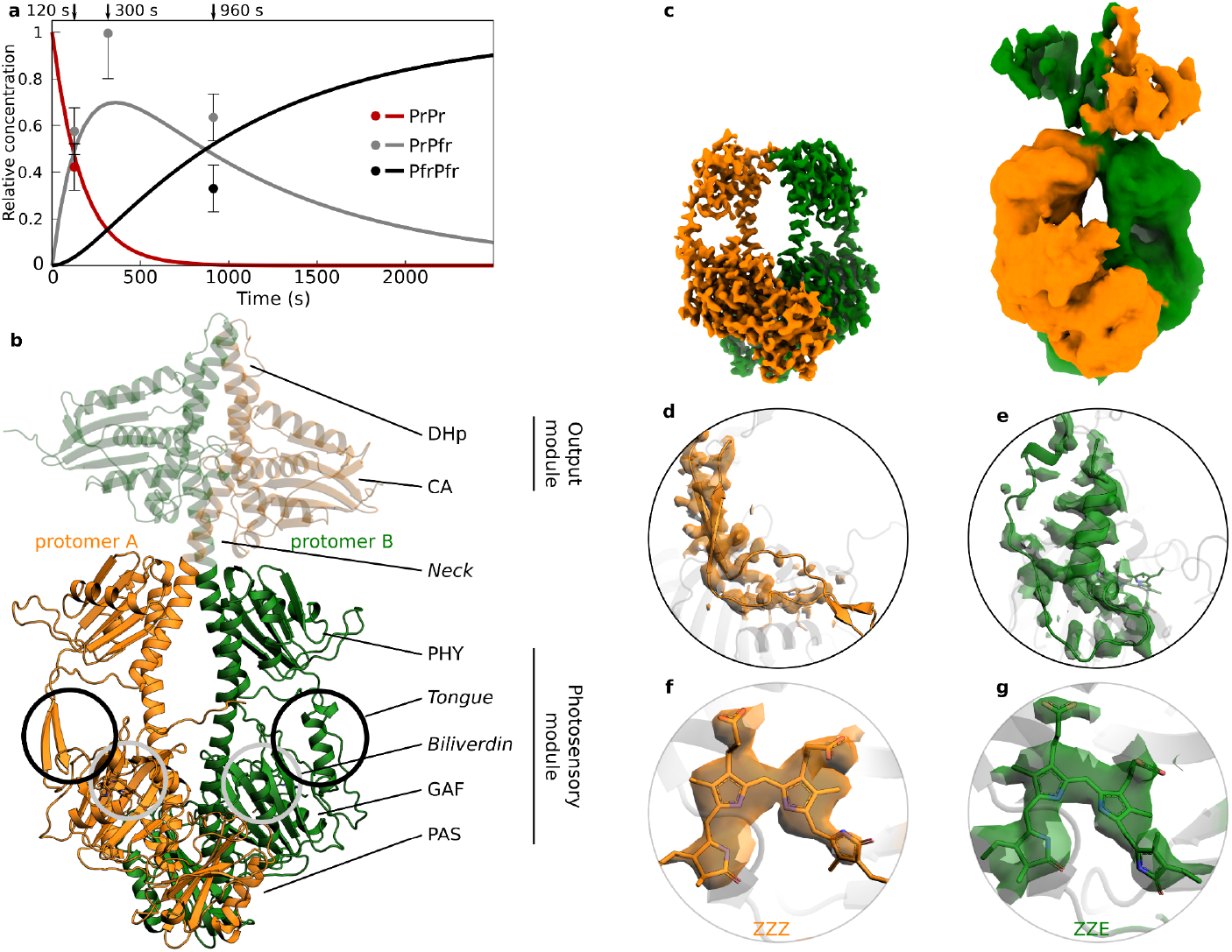
The dark reversion intermediate of PaBphP is a PrPfr hybrid. (a) The presence of a PrPfr intermediate state in the dark reversion reaction of PaBphP was revealed by UV-Vis absorbance spectroscopy at 20°C.^38^ Here, the time-dependent concentrations based on the rates identified by Prodhan et al. (2025) are shown. The time points at which cryo-EM data were recorded are indicated. The dots show the relative concentrations estimated from the number of particles assigned to each 3D volume in the reconstructions; the error bars are set to 0.1. At 300s the uncertainty was higher and the error bar for the PrPfr state was set to 0.2. PfrPfr particles were observed as 2D classes, and likely some PrPr particles are included in the 3D volume for the hybrid state, but neither of the homodimer states could be refined. (b) The cryo-EM structure of the PrPfr hybrid, reconstructed from the 300 s dark reversion data (the output module, not clearly resolved in the reconstruction, is not included in the final model but is shown here with decreased opacity). (c) The panel shows the coulomb density reconstructed via heterogeneous reconstruction of the full-length protein (right) and local refinement of the PSM (left). (d-g) The PHY-tongue (d, e) and the chromophore (f, g) show characteristic conformations for Pr-state in protomer a (d,f) and Pfr-state in protomer b (e,g).

In the absence of light, phytochromes return to their resting states. Most phytochromes dark-revert to the Pr state, although “bathy” bacterial phytochromes revert to Pfr.^28^ The dark reversion rates are species-dependent and modulated by pH, phosphorylation, and protein-protein interactions.^29–32^ Dark reversion plays a key functional role by determining the lifetime of the photoactivated state and thus tuning the organism’s sensitivity to ambient light cues. Evolution of paralogous genes allows plants to host several isoforms of phytochromes with different dark reversion rates, which, despite having partially similar spectral properties, can cater for a large dynamic range of signalling.^33^ Additionally, since the reversion rate is highly temperature-dependent, phytochromes can act as biological thermometers.^6,34,35^ Thus, the protein must be able to control the thermal reversion reaction of the bilin chromophore, but the structural and chemical basis for this remains poorly understood.

Phytochromes are typically dimeric proteins. This significantly enhances the signaling capabilities of the proteins, exemplified by the formation of PrPfr heterodimers, creating a distinct “third state” in addition to the PrPr and PfrPfr homodimers.^5,24,36^ In *Arabidopsis thaliana* PhyB the PrPfr heterodimer undergoes dark reversion approximately 100 times faster than the PfrPfr homodimer, which indicates strong intra-dimeric allostery and leads to effective suppression of PhyB-mediated signaling under low light intensity.^37^ While the existence of heterodimeric states has long been recognized in plants, evidence for PrPfr states in bacterial phytochromes is scarce. To date, only one crystal structure of an engineered bacterial phytochrome and a recent cryo-EM structure from a myxobacterial phytochrome have captured such hybrid forms.^20,24^ In our recent work using optical spectroscopy, we show that PrPfr heterodimeric states are indeed populated during dark reversion in bacterial phytochromes.^38^ Notably, we observed clear evidence of allosteric regulation of the dark reversion rate.

Despite its physiological importance, dark reversion remains poorly understood at the structural and chemical levels. For thermal reversion, the chromophore has to isomerize around the 15-16 double bond (**Fig. S1**), which is expected to have a substantial activation energy on the order of 100 kJ/mol.^39^ This corresponds to about 40 times the kT at 298 K, and thus prohibitively slow rates would be expected. However, faster rates are observed, implying that chemical mechanisms must be at play to lower the activation energy. One proposed mechanism that fulfils this requirement is the keto-enol tautomerization on the D-ring of the biliverdin in its Pr form, which reduces the bond order of the C15-C16 bond and hence lowers the activation barrier for thermal reversion.^40–42^ In this model, the enol form has to be formed before Z to E isomerization of the biliverdin can occur. Quantum chemical calculations suggest that this is possible when water molecules participate in the transition state between keto and enol forms.^41^ Moreover, pH-dependent spectroscopy suggests that the concentration of the enol state correlates with the protonation states of a histidine in the vicinity of the chromophore.^43^

From these considerations, the question arises as to how phytochromes exert control over the dark reversion reaction, both in terms of inter-dimer allostery and with respect to the dark reversion mechanism itself. A realistic candidate is conformational selection, where two or more substates co-exist in the protein, with differential propensity for dark reversion. Detection of the structure of such substates and particularly those with low abundance is a major challenge.^44^ Crystal structures of phytochromes are unlikely to capture this state, because the active substates are frozen out at low temperatures and in the crystal lattice. NMR and spectroscopic studies have indicated the presence of multiple ground state conformations of phytochromes in solution,^45–48^ but lack structural specificity. Thus, the conformational selection mechanism remains unproven, and a structural understanding of how the dark reversion reaction is controlled is still lacking.

Here, we use single-particle cryo-EM to solve the structure of the bathy phytochrome from *Pseudomonas aeruginosa* at several time points during the dark reversion. Intriguingly, we observe that the dark reversion is strongly asymmetric, where only one protomer reverts to form a hybrid PrPfr state. This raises the opportunity for identifying the structural mechanisms with which the thermal isomerization of the biliverdin is controlled from within the same phytochrome.

## Results

### PrPfr heterodimer is confirmed as dark reversion intermediate

In order to solve the structure of the PrPfr heterodimer state during dark reversion of *Pa*BphP, we made use of its expected relative concentration as derived from spectroscopic observation (**Fig. 1a**),^38^ and cryo-trapped the protein on electron microscopy grids at 120 s, 300 s and 960 s after photoexcitation with a laser diode at 780nm. At 120 s, a new structural state (**Fig. 1b, c**) is observed in the protein ensemble (approximately 58% of the particles) along the known PrPr structure (see **Fig. S2** for details of the reconstruction).^25^ At 300 s, the new state dominates the ensemble on the grids. Neither the homodimeric PrPr nor the PfrPfr states could be separately reconstructed, but a small number of open, PfrPfr-like particles can be identified during 2D classification (**Fig. S3**). At 960 s, a mixture of the new state (approximately 64% of all particles) and the open PfrPfr state (approximately 36%) is present (**Fig. S4**).

The newly observed structural state is a PrPfr heterodimer (**Fig. 1b**) with one protomer in Pr and the other one in Pfr, as judged based on the secondary structure of the PHY-tongue (**Fig. 1d, e**), isomerization state and location of the chromophore (**Fig. 1f, g**), and the rotamerization states of key residues lining the binding pocket (**Fig. S5**). The structure of the PrPfr heterodimer was highly similar at all three time points and was, next to the already known PrPr and PfrPfr states, the only structural state identified from the particle ensemble at any of the time points.^25^ It is difficult to determine the fractional population of the three states quantitatively from the analysis of the cryo-EM data, however, the estimated concentrations agree well with the kinetic prediction (**Fig. 1a**). This confirms that the spectroscopic intermediate indeed is a PrPfr heterodimer.

### Two geometries at the dimer interface resolved for PrPfr and PrPr states

The observed PrPfr heterodimer structure is naturally asymmetric at the tongue and chromophore regions, due to the different photochromic states. However, we also observe strong asymmetry beyond these regions at the dimer interface of the PAS/GAF domains. Here, two distinct geometries are observed. In geometry I (**Fig. 2d**), consisting of the GAF_A_ and PAS_B_ domains (subscripts denoting the protomer), helix A_A_ is in a conformation consistent with the Pfr state crystal structure, making contact with spine_B_, but not connecting to PAS_B_. This results in an open interface (**Fig. 2b and d**).^17^ Geometry II (**Fig. 2e**) involves GAF_B_ and PAS_A_ and here helixA_B_ is oriented into a 3-helix bundle with helix B_B_, spine_B_, and makes direct contact with PAS_A_. This results in a closed interface (**Fig. 2b and e**). Highly similar geometries are also observed for the PrPr state and a characteristic gap between GAF_A_ and PAS_B_ (see arrow) is observed in both PrPr and PfrPr (**Fig. 2a**).^25^ **Fig. 2c-e** visualize how the interaction network of helix A changes between the two interface geometries. Several key interactions change, affecting the neighbouring spine helix, helix B, and the connection to residues in the PAS domain. We conclude that the interface region at the PAS-GAF domains is intrinsically asymmetric in PrPr and PrPfr (but not in PfrPfr)^25^ and that the positioning of helix A plays a decisive role in breaking the symmetry. During reconstruction of the cryo-EM maps, this asymmetric dimer arrangement drives alignment, resulting in high-confidence assignments for protomers A and B in both the PrPr and PrPfr states.

**Fig. 2.**
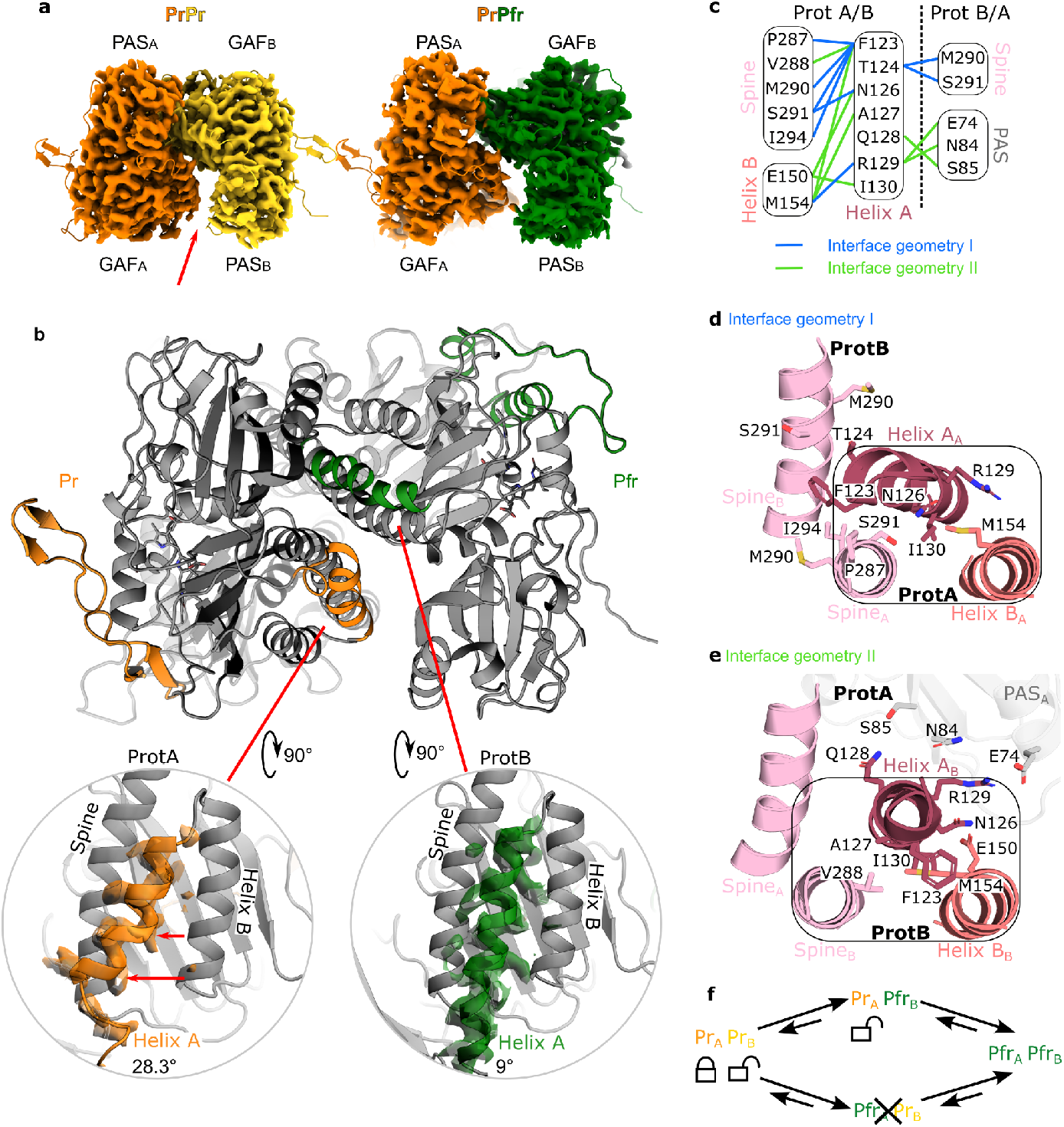
The PSM dimer interface is asymmetric in both the PrPr ground state and the PrPfr intermediate. (a) Two distinct PAS-GAF interaction geometries are observed in the PrPr and the hybrid PrPfr states, conserved during the transition from one to the other (red arrow points at the gap related to geometry I). (b) The orientation of helix A differs between the two protomers and correlates strongly with the photochromic state of the protomer in PrPfr: the relaxed helix A, the α-helical PHY tongue and ZZE state chromophore are all exclusive to protomer B (green). This implies a strongly asymmetric dark reversion reaction, in which only protomer B dark converts from PrPr to PrPfr. The angle in the insets is calculated between helix A (residues 123-136) and the base of the spine helix (residues 286-300). (c) Distinct interface geometries according to interactions (interaction frequency differences) for each Helix A of PrPr are shown schematically. Geometry I (blue) shows interactions of ProtB and geometry II (green) shows interactions of ProtA. Structural representations highlight the shift from interactions of Helix A_A_ to Spine_B_ (d) to interactions of Helix A_B_ with PAS_A_ (e). (f) Schematic of the observed dark reversion sequence: the ProtA-Pfr hybrid state was not found in the cryo-EM data and was determined to occur less frequently in the MD simulations.

### Only one protomer of the asymmetric dimer dark-reverts from PrPr to the PrPfr hybrid state

Since the asymmetric arrangement of the PAS/GAF dimer interface is retained between PrPr and PrPfr, it provides a handle to trace protomers A and B separately in their dark reversion process. If the dark reversion was stochastic between the protomers, either protomer A or B could backconvert, and blurred densities would be expected at the chromophore and tongue positions because the two conformations would be averaged over each other. However, this is not observed. Instead, we recover high-resolution information at the chromophore and tongue in both protomers, which allows clear assignment to each one of the photochromic states (**Fig. 1 d-g**). This leads to the surprising observation that only one of the protomers (protomer B) dark-reverts from the PrPr to the hybrid PrPfr state (**Fig 2f**). We conclude that the dark reversion reaction in protomer A must be very slow or inhibited. It seems very likely that the difference in reactivity is caused by the observed structural asymmetry of the dimer in PrPr.

### A switch of H277 and its hydrogen bonding to the chromophore explains how dark reversion is controlled

Control over dark reversion requires regulation of the isomerization reaction at the biliverdin. In order to identify this mechanism, we reanalyzed our previously published electron density map (EMD-19981) and model (PDB 9EUT) of the PrPr state.^25^ **Fig. 3a** and **c** show that the density around the biliverdin in protomer B is weaker than in protomer A, which indicates a higher degree of heterogeneity in protomer B compared to A. Moreover, there is a difference in the conformation of the conserved histidine 277 (H277) with respect to the biliverdin: in protomer A the distance of the carbonyl oxygen of ring D of the BV (OD) to the epsilon nitrogen on the histidine (NE) is 2.6 Å with an angle of 150°, which is a suitable hydrogen bonding geometry; while in protomer B this distance is increased to 2.9 Å with an angle of 110°, which is not well compatible with hydrogen bonding in protomer B. Local resolution of the maps around the chromophore is ∼2.4 Å, which allows for confident placement of the histidine side chains.^25^ Additionally, the maps provide clear support for the refined positions of H277 as its electron density extends further towards the BV in protomer A compared to B **(Fig. 3a** and **c**). Further down we will extend the structural analysis and find that the cryo-EM densities likely represent a mixture of substates. It is intriguing that the structural observations correlate with the ability of the protomers to react back to Pfr and suggests that, when hydrogen bonding of H277 to the D-ring carbonyl is direct as in protomer A, the dark reversion reaction becomes inhibited.

**Fig. 3.**
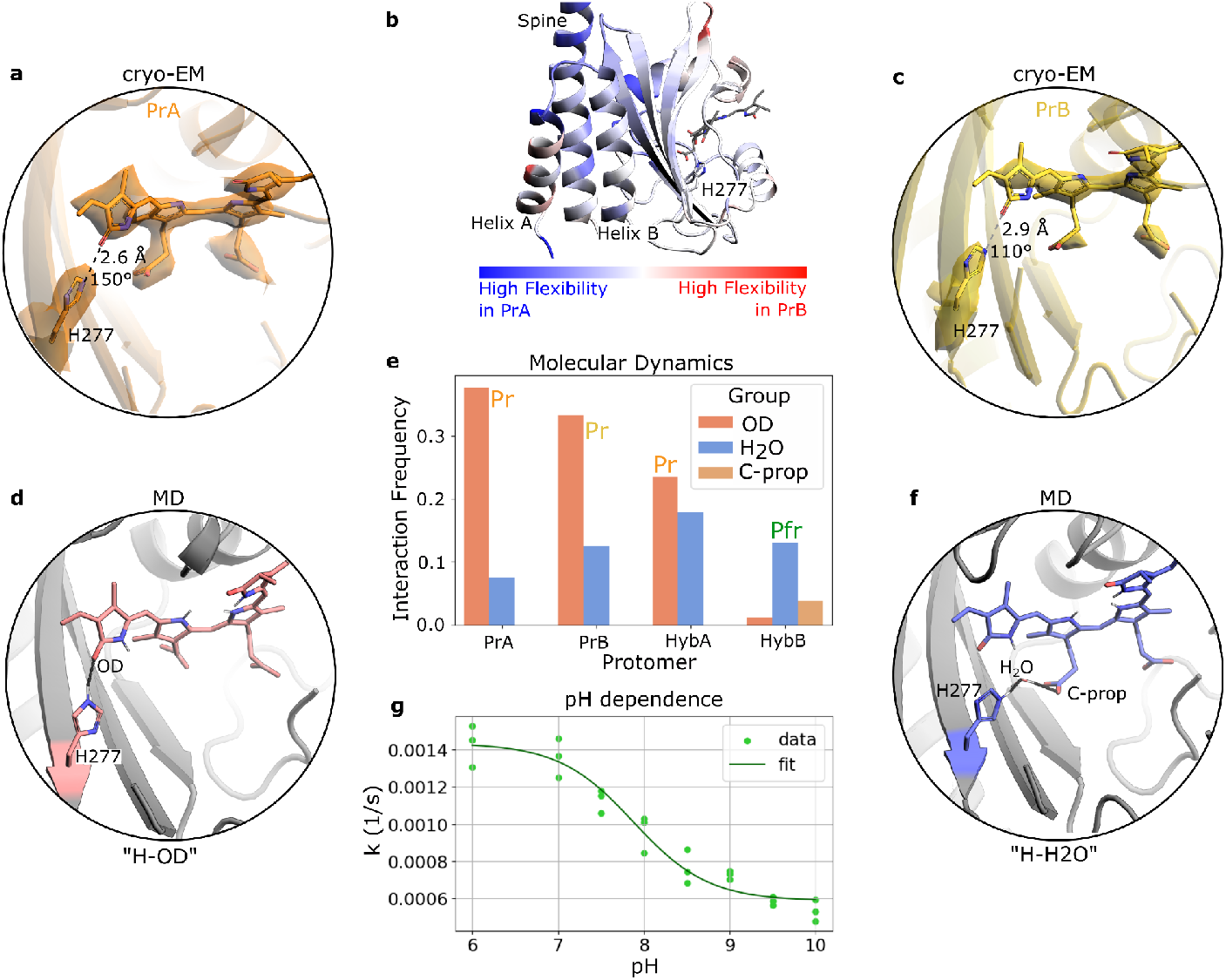
Distinct interaction pattern of H277 with BV reveals back-reversion advantage of protomer B. (a, c) Densities of biliverdin at identical contour levels in protomers of PrPr with distances from H277-NE to BV-OD and angles from H277-NE via H277-H_NE_ to BV-OD. (b) The difference in RMSF of the two protomers (which is comparable to an experimental B-factor and a measure for residue-wise flexibility) and highlights a rigidification in protomer B around the active site. (e) Polar contact (hydrogen bonds) frequency of the different protomers where H277 interacts with biliverdin (H277-OD, “H-OD”) directly or via bridging waters (H277-H2O-Cprop, “H-H2O”) in each protomer of PrPr (PrA and PrB) and PrPfr (HybA and HybB). (d, f) Structural evaluation of the different states of the binding pocket: (d) H277-OD and (f) H277-H2O-Cprop. OD is the carbonyl of the D-ring and C-prop describes the carboxylates of the C-propionate. (g) pH dependence of dark reversion kinetics (rate constant vs pH) shows pKa of 7.9. For additional details, see Fig. S9.

To further elucidate the dynamics of the biliverdin and H277, we performed three repetitions of molecular dynamics simulations (MD) for both the PrPr and PrPfr states. Within the 250 ns simulation time, the interface geometries predominantly stayed constant (**Fig. S6**). Using these trajectories, we performed an interaction analysis, where the frequency of contacts between residues is determined according to geometric and electrostatic cutoffs. Higher interaction frequencies thereby indicate a higher stability of the state. We find two predominant conformations, where H277 either forms a hydrogen bond directly to the D-ring (pattern “H-OD”, **Fig. 3d** and **S1**) or it forms a bond via a water molecule to the propionate group of ring C (pattern “H-H2O”, **Fig. 3f**). The switch is manifested by retracting and twisting movement of the imidazole ring of H277 which leads to the change of hydrogen bonding. The fact that the trajectories sample both conformations in all of the MD trajectories implies that the states are close in energy and that they can interconvert within nanoseconds (**Fig. S7**). The switch in hydrogen bonding of H277 has a direct effect on the conformation of the D-ring, which twists around V5 (torsion between C and D ring, NC-C14-C15-C16, see numbering in **S1**) by 7.1° (**Fig. S8**).

**Fig. 3e** summarizes the frequencies with which each of the two predominant conformations occurs in the simulations. For the PrPr state, we find that the “H-OD” pattern is prevalent in protomer A (“H-OD” ∼38 %, “H-H2O” ∼7 %). In protomer B, the occurrence of the “H-H2O” increases while “H-OD” decreases (“H-OD” ∼33 %, “H-H2O” ∼12 %). This suggests that hydrogen bonding between H277 and the D-ring is tighter in the non-reactive protomer A. While the absolute frequencies of these patterns from MD trajectories may not be 100% reliable because of computational limitations (i.e. imperfect force field and water model), the increased “H–H2O” interactions observed between the two otherwise highly similar protomers support the robustness of the observed trend. The observation suggests that the regulation of the dark reversion reaction occurs by control of the abundance of the reversion-ready “H-H2O” conformation, where H277 is not bound to the D-ring. Importantly, this conclusion aligns with the observations from cryo-EM maps (**Fig. 3a** and **c**).

Turning to the MD simulations of the hybrid PrPfr state, we intriguingly find that the “H-H2O” conformation of H277 in protomer A increases in frequency compared to PrPr (“H-OD” ∼23 %, “H-H2O” ∼18 %). This finding is supported by the cryo-EM structures of the hybrid state, where the H277 in protomer A of PrPfr is refined into a position with distance of 2.6 Å and 120° with respect to the D-ring carbonyl, which is a conformation that resembles that of protomer B in PrPr and is unfavourable for direct hydrogen bonding. This suggests that protomer A can now undergo dark reversion from PrPfr to PfrPfr (**Fig. 3e**).

### pH-dependent dark reversion rates support the role of a histidine residue in controlling chromophore isomerization

To experimentally probe the proposed role of the histidine residue in the dark reversion process, we measured the pH dependence of the dark reaction kinetics (**Fig. 3g** and **Fig S9** for additional information). The data revealed a sigmoidal decrease in rate constants with rising pH between values 6 and 9, with an apparent transition midpoint near pH 7.9. The type of pH profile indicates that a single titratable group controls the reaction kinetics. These results are consistent with H277 regulating biliverdin isomerization and support the interpretation of the cryo-EM data and MD simulations.

### Protein dynamics link the H277 switch to conformation of the dimer interface

After highlighting the importance of the positioning of H277 for dark reversion (**Fig. 3**) it is further relevant to elucidate how the asymmetry at the dimer interface and in particular the positioning of helix A (**Fig. 2**) are connected to this molecular switch. Inspection of the MD trajectories and the cryo-EM structures did not reveal a clear pathway of concerted structural changes. Instead, we find correlated differences in the dynamics as modelled by the MD simulations. In protomer A, the spine helix, helix B, and the five-membered beta-sheet of the PAS domain, which connects to H277, are notably more flexible than in protomer B according to the RMSF (**Fig. 3b**). Only helix A is more rigid in protomer A pointing at a stabilization due to the formed interactions with spine_B_. The observation suggests that the asymmetric arrangement of helix A at the interface (geometry I, **Fig. 2d**) and the reactivity of the biliverdin are connected via altered dynamics of the PAS/GAF domains. Rigidification of the active site enables stabilization of the surrounding residues and thus opens up the space for H277 to retract from the direct hydrogen bonding with biliverdin.

In conclusion, we find two patterns of hydrogen bonding of H277, which correlate with the dark reversion propensities of the two protomers. This leads to strongly asymmetric dark reversion: Thermal isomerization of the biliverdin in protomer A is blocked or prohibitively slow in PrPr, which corresponds to the dominant occurrence of hydrogen bonding of H277 directly to the D-ring. However, in protomer B, the biliverdin is not hydrogen bonded to H277 and the reversion reaction occurs. In the PrPfr heterodimer state, the hydrogen bond pattern of protomer A is altered, with H277 binding much more frequently in the “H-H2O” arrangement, thus freeing up the biliverdin for thermal isomerization. The cryo-EM structures show that this molecular switch of H277 is correlated with the asymmetric structure at the dimer interface and the MD simulations suggest that the signal is transduced by modulation of the internal dynamics in the PAS-GAF domains.

## Discussion

### The PrPfr intermediate solved from dark reversion may be functionally important

PrPfr intermediates in the phytochrome photocycle have been described in plants and, more recently, in bacterial systems.^5,20,24,36,37^ Here, we demonstrate that the PrPfr state is a key intermediate in the dark reversion reaction of bacterial phytochromes, structurally confirming our recent spectroscopic assignment.^38^ In plants, it is established that the intermediates are functionally relevant, enhancing the signalling potential of the protein.^37^ Thus, even bacterial phytochromes may use the PrPfr hybrid states for signalling. The role of *Pa*BphP in its organism is to initiate biofilm formation in darkness when the bacteria enter the body.^49^ This is consistent with the comparatively fast dark reversion time, but it is not obvious how allosteric regulation or the presence of a hybrid state would be beneficial for this. We note, however, that state-of-the-art functional assays have not explicitly considered the possibility of a transient hybrid state for *Pa*BphP and other bacterial phytochromes, thus it could be that the function of the PrPfr hybrids have so far eluded detection.^50,51^

### The PAS-GAF dimer interface is an important and previously unrecognized regulatory site

A large number of structural elements, including the chromophore, its direct environment, the PHY-tongue, and the output domains have been implicated in the photo- and dark reactions of phytochromes.^15–20,42,52^ In this work we add the dimer interface at the PAS-GAF domains to this group and find that the distinct conformations of helix A in the two protomers strongly correlates with the ability of the connected chromophore to undergo dark isomerization. Notably, photoinduced changes in the interface region were also observed in an NMR investigation of a monomeric variant of the *D. radiodurans* phytochrome and in the cryo-EM structures of the PrPr, PrPfr, and PfrPfr states of the canonical bacteriophytochrome *Sa*BphP2.^24,53^ In the latter study, the PrPfr heterodimer had an asymmetric position of helix A (**Fig. S10**) and it is conceivable that this allosterically controls the rate constants of reversion in *Sa*BphP2, in analogy to the findings made here for *Pa*BphP.

### The histidine switch works together with keto-enol tautomerization as a key regulator in the dark reversion process

Importantly, our structural data suggest a conformational selection mechanism for phytochrome dark reversion, in which substates of H277 are in equilibrium and their relative abundance regulates the reaction kinetics (**Fig. 4**). In the PrPr state, H277 adopts two distinct conformations. In the first one it forms a direct hydrogen bond with the D-ring carbonyl, whereas in the other one it connects to the C-propionate group via a water molecule (**Fig. 4b**). The relative abundance of the two states is allosterically controlled by the symmetry break at the dimer interface (**Fig. 4c**) and dynamics in the PAS/GAF domains link the dimer interface and H277 (**Fig. 4a**). These changes are small in amplitude, but are large enough to alter the hydrogen bonding environment of the D-ring, thereby resulting in large effects on dark reversion rates.

**Fig. 4.**
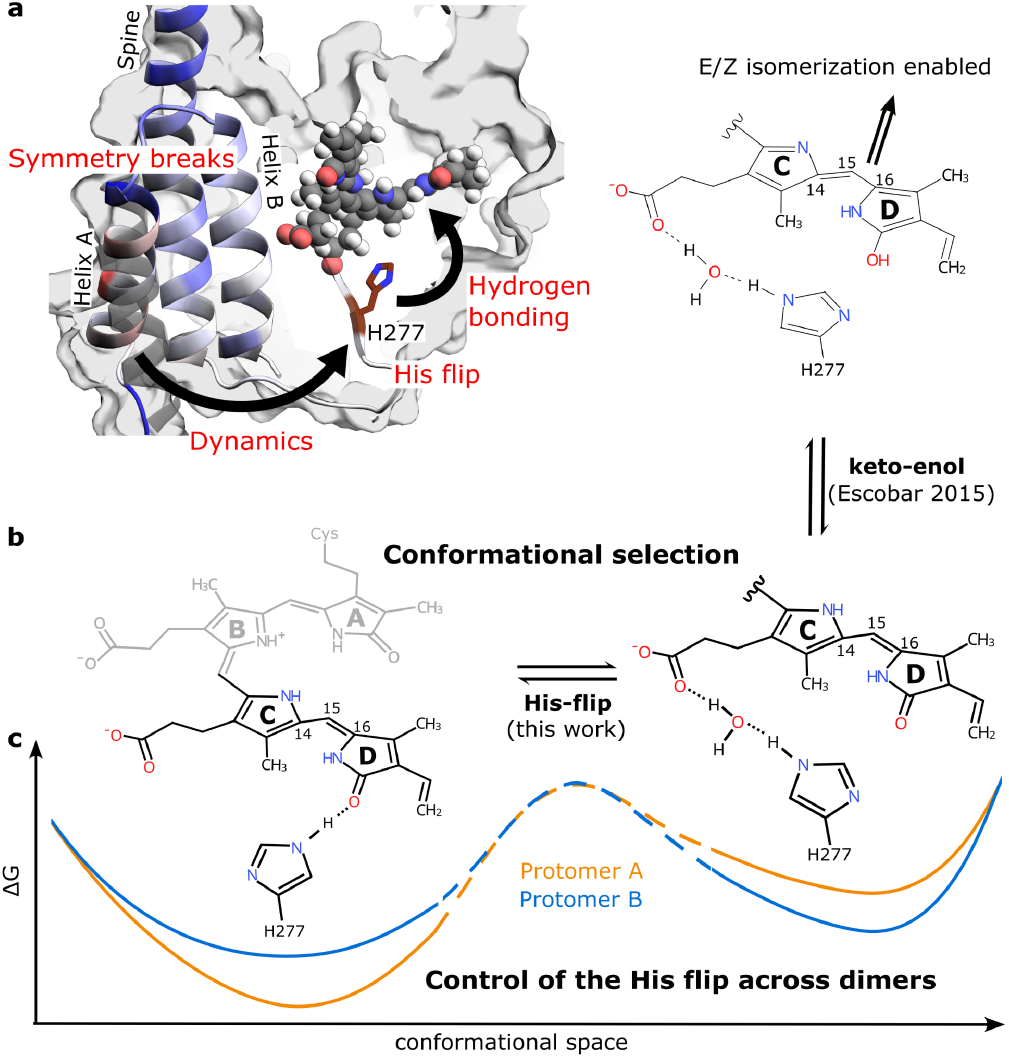
Summary of the asymmetric dark reversion mechanism. (a) Close-up of a phytochrome PAS-GAF domain shows the influence of the asymmetric interface (Helix A) on H277 (dynamics) and BV (hydrogen bonding). The helices are colored according to the difference in RMSF of the two protomers (blue means protomer A is more flexible) and the alternative conformation of helix A is shown in grey. (b) Combining the keto-enol tautomerization idea by Escobar et. al.^42^ with the difference in hydrogen bonding of H277 described in this work and highlighting the importance of conformational selection for the mechanism. (c) The schematic free energy landscapes correspond to the observed shifts in abundance of the different sub-states. Protomer A prefers the “H-OD” conformation (left) and protomer B the “H-H2O” conformation (right) when in Pr. Considering the higher occurrence of “H-H2O” in protomer B, from which dark reversion reaction can occur, this is in line with protomer B dark reverting first, coherent with the observations from the cryo-EM structure of the PrPfr hybrid state.

The switch of H277 identified here, works in concert with the suggested keto–enol tautomerization mechanism of dark reversion in bathy phytochromes.^42^ In this mechanism, internal proton transfer from the B/C-rings to the D-ring carbonyl reduces the double-bond character between C15 and C16, thereby lowering the barrier for ring rotation. When H277 binds directly to the D-ring carbonyl, it stabilizes the keto form and inhibits tautomerization into the enol form (**Fig. 4b**), thereby effectively blocking isomerization. When H277 switches to its alternative conformation and binds to the C-ring propionate via a water molecule, the carbonyl is freed from hydrogen bonding, may transition into the enol form, and dark reversion can proceed. Our (**Fig. 3g**) and previously measured pH dependencies of the dark reversion further confirm the balance-tipping role of a histidine for the dark reversion kinetics.^42,54^ Moreover, the proposed mechanism is in line with quantum chemical computations of the keto-enol reaction, which emphasize the importance of water molecules for access to the enol as they stabilize the transition state between the two forms.^41^ This is fulfilled for the conformation where H277 does not bind to the D-ring C=O.

### Cryo-EM is a promising technique for measuring conformational heterogeneity

As a single-molecule technique, cryo-EM is in principle well suited to measure conformational heterogeneity such as detected here. However, it still struggles to reach the resolution needed to resolve chemically relevant differences in small and/or flexible complexes, as the conformational changes we detect in our system are too small in amplitude to be sorted from the data. The detection of low-abundance, high-energy states has so far been driven mainly by NMR, but our work shows the way for cryo-EM to achieve the same.^44,55^ In the present case, we had to complement the structural data with molecular dynamics simulations, but future methodological advances may enable cryo-EM to characterize such states more autonomously.

## Conclusion

Here, we have shown that the conserved H277 and the surrounding hydrogen-bonding network play a key role in regulating the dark reversion reaction of phytochromes. H277 is widely conserved across the phytochrome family and has been shown to alter the isomerization yield of the Pr-to-Pfr photoreaction.^28,56–58^ The mechanism is also in general agreement with the detection of structural heterogeneity for phytochromes from NMR and time-resolved optical spectroscopy experiments.^45–48^ Allosteric control of dark reversion rates is a common feature of phytochromes.^36,59^ Here, we show that a break in symmetry at the dimer interface, linked to a change in hydrogen bonding of H277, is a key chemical factor controlling this process. We hope that this conformational selection mechanism of the isomerization reaction is of use for the design of future optogenetic phytochrome variants.

## Methods

### Protein purification

*Pa*BphP was recombinantly expressed in *E. coli* BL21 cells transformed with the *Pa*BphP gene (GenBank: AAG07504.1, on a pET28a(+) backbone) and a heme oxygenase gene was kindly provided by Prof. Janne A. Ihalainen. The cells were grown in LB media containing 50 μg/mL kanamycin and 34.5 μg/mL chloramphenicol at 37°C until OD_600_ reached 0.6, then supplemented with 1 mM IPTG and 1 mM 5-aminolevulinic acid before lowering the incubation temperature to 18°C.

The cells were processed in lysis buffer (50 mM Tris, 150 mM NaCl, 10% glycerol, 10 U/mL of DNase 1 and one tablet of cOmplete EDTA-free protease inhibitor (Roche) at pH 8.0) using an EmulsiFlex C3 homogeniser (Avestin), and the lysate briefly incubated with a molar excess of biliverdin hydrochloride. The protein was then purified through a two-step chromatography protocol: first, Ni-NTA affinity chromatography (HisTrap HP 5 mL, Cytiva) by washing the bound sample with wash buffer (50 mM Tris and 1 M NaCl at pH 8.0) and eluting with elution buffer (50 mM Tris, 50 mM NaCl, 300 mM imidazole at pH 8.0); followed by size-exclusion chromatography (HiLoad 16/600 Superdex 200 pg, Cytiva) in SEC buffer (30 mM Tris at pH 8.0). The purified protein was collected at a concentration of 3 mg/mL and flash-frozen for storage.

### Cryo-electron microscopy specimen preparation

For structural studies, the protein was diluted to 1.5 mg/mL and the buffer exchanged to 80 mM Tris, 10 mM MgCl2 and 150 mM CH3CO2K at pH of 7.8. The bulk sample was illuminated with a 780 nm light source over 30 s (total dose: ∼1 mJ/mm^2^), then left in complete darkness until deposition on glow discharged UltrAuFoil (Quantifoil) R 1.2/1.3 (300 mesh) grids and flash-freezing in liquid ethane. The dark reversion time was measured from the removal of the far-red light source and was considered terminated at the moment of flash freezing. Specimens with 120, 300 and 960 seconds of dark reversion were prepared. The photon dose is sufficient to achieve full conversion into Pr.

### Cryo-electron microscopy data collection

The specimens were imaged using a Titan Krios G2 (Thermo Scientific) 300 keV transmission electron microscope equipped with a K3 (Gatan) detector and a BioQuantum (Gatan) energy filter. The data acquisition settings were as follows: 0.828 Å/px (105kx magnification), 50 e/Å^2^ total electron dose, -0.8 to -2.0 mm defocus range, 20 eV energy filter slit width. Automated collection was performed using the EPU software (Thermo Scientific). A total of 13,534 movies were collected for the 120 s time point, 14,573 movies for the 300 s time point, and 13,112 movies for the 960 s time point.

### Cryo-electron microscopy data processing

The collected micrographs were processed using Cryosparc v4.6.2. Preprocessing consisted of patch motion correction, patch CTF estimation and micrograph denoising (only for visualization purposes). A total of 13,393, 14,033, and 10,323 high-quality micrographs were accepted based on CTF fit, average defocus and relative ice thickness. Particles were originally picked as 80-180 Å blobs. For the 300 s dataset, a Topaz model was trained for more accurate picking, whereas in the other datasets the originally picked particle sets were used for further refinement. Removal of junk particles was done by subsequent rounds of 2D classification and heterogeneous refinement using both the open and closed dimers and several “junk” volumes as references.

For the dataset at 300 s delay, a total of 964,467 particles were identified, which contributed to a PrPfr reconstruction. Further 3D classification did not reveal a separate PrPr class, but it was used to further select 488,190 particles with especially well-ordered α-helices in the PHY_B_ region (**Fig. S3**). The failure to achieve a PrPr reconstruction is not proof of its absence in the population, but indicative of bias towards the more populated PrPfr state during reconstruction. To account for this increased uncertainty at this time point, we increased the error estimate to 0.2 in **Fig. 1**. With the 488,190 particle subset, separate local refinements were conducted with masks for the PAS-GAF-PHY regions of protomers A and B, which yielded maps of global resolution 2.87 and 2.85 Å, respectively. The output module was resolved at a nominal resolution of 5.4 Å. An atomic model was built using the ISOLDE (v1.6) extension of ChimeraX (v1.8), with the models 9EUY and 9EUT used as templates.^25,60^ The two locally refined maps were then combined using the Phenix function ‘Combine focused maps’.

For the dataset at 120 s delay, a total of 895,611 particles were found. These were subjected to 3D classification using the “simple” initialization mode to reveal reconstructions of both the PrPr and the PrPfr states, followed by a second, targeted 3D classification round. 376,742 (42.1%) of the particles contributed to the PrPr map, while 513,575 particles (57.9%) yielded a PrPfr volume (Fig. S2). Local refinements of the PAS-GAF-PHY regions (excluding the output module) were conducted on both classes, resulting in global resolutions of 3.02 and 2.82 Å, respectively. Comparing the two maps against 9EUT and the newly built PrPfr models showed good fits, so no new model was built into these refinements.

For the dataset at 960 s delay, a total of 476,372 particles were identified, with 303,375 (63.7%) contributing to a PrPfr map at 3.07 Å resolution, and 172,997 (36.3%) particles belonging to an open PfrPfr reconstruction, resolved at 3.29 Å resolution (**Fig. S4**).

Estimation of relative concentrations at each time point was done by generating a consensus reconstruction (where applicable), then splitting the final particle set into Pr/PrPfr/Pfr groups by 3D classification in input initialization mode, supplying high-resolution (<3 Å) input maps and filtering to 4 Å.

### Spectroscopy

UV-Vis spectroscopic measurements for the pH-dependent dark reversion rates were conducted under conditions as described by Prodhan et al. (2025): the protein was kept in a buffer consisting of 50 mM Tris and 50 mM NaCl at a concentration of 0.6 mg/ml, with pH ranging from 6.0 to 10.0.^38^ The temperature was set to 25°C. The sample was kept in complete darkness or under dim green light during preparation, and at the start of the measurement was illuminated with a 780 nm light source over 30 s (total dose: ∼1 mJ/mm^2^). The dark reversion was followed by collecting full UV-Vis spectra (250-800 nm) at 45 s intervals with a total of 100 repeats using a Shimadzu UV-1900i spectrophotometer equipped with LabSolutions UVVis software (Version 1.13). The first spectrum was collected 5 s after far-red illumination. The absolute decay of Pr (at 698 nm) and growth of Pfr (at 749 nm) was fitted with a general exponential function:

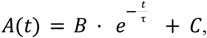

With B, C and τ as parameters to fit. The rate constant k is the reciprocal of τ (k = ^™1^). We then fitted the k vs pH plot with a sigmoidal function, i.e. a Henderson-Hasselbalch-like function:

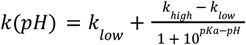

Where k_low_ is the rate constant at low pH and k_high_ is the rate constant at high pH. From that the pKa was directly extracted.

### Molecular dynamics simulations

We performed molecular dynamics simulations of both the full length PrPr and hybrid PrPfr states. Thus, the structures were protonated with H++ at pH 8 according to the experimental conditions (H277 is protonated as HIE).^61–64^ As force-field ff14SB^65^ was used as well as the TIP3P^66^ water model (10 Å minimum wall distance),^67,68^ utilizing the AMBER24 tool kit, i.e. the tleap implementation.^69^ Charges were neutralized with a uniform background charge and the systems were equilibrated with a multi-step protocol.^65,70,71^

Both systems were simulated for approximately 250 ns three times (see **Table 1**) in an NpT ensemble (using pmemd.cuda),^72^ using a timestep of 2 fs, since the SHAKE algorithm was applied.^72,73^ The particle-mesh Ewald (PME) method took care of electrostatic interactions. The langevin thermostat kept the temperature at 300 K and the monte carlo barostat kept pressure at 1 bar.^74,75^

**Table 1.** Summary of simulation frames and time of all repetitions.

| Simulation repetitions | PrPr |  | PrPfr |  |
| --- | --- | --- | --- | --- |
|  | Frames | Time | Frames | Time |
| 1 | 12465 | 249 ns | 11901 | 238 ns |
| 2 | 17832 | 357 ns | 12500 | 250 ns |
| 3 | 16472 | 329 ns | 15271 | 305 ns |

The performed simulations were analysed and visualised with cpptraj, vmd and PyMOL (The PyMOL Molecular Graphics System, Version 3.0, Schrödinger LLC.).^76,77^ RMSF (Root mean square fluctuation) calculations and interaction analysis were performed with cpptraj and the GetContacts tool (https://getcontacts.github.io/) and an associated python script.^78^ For the RMSF the structures were aligned on the respective GAF domain before calculation of the RMSF which were thus subtracted from one another (scale ranges between 2 Å). To show that the assignment is reliable we calculated the difference of each protomer A to each protomer B across all PrPr simulations (**Fig. S11**). For the hydrogen bond analysis of H277 we used the cpptraj command hbond and selected as donor H277 and as acceptor the chromophore. For the interface interaction analysis, we combined hydrogen bonds, salt bridges and van der Waals interactions calculated with GetContacts and applied a cutoff of 0.5 for the difference between the interaction frequencies to highlight the main differences between the two geometries.

## Acknowledgements

We acknowledge the use of the Cryo-EM Uppsala facility for cryo-EM specimen preparation and screening, funded by the Department of Cell and Molecular Biology, the Disciplinary Domains of Science and Technology and of Medicine and Pharmacy at Uppsala University.

The cryo-EM data was collected at the Cryo-EM Swedish National Facility funded by the Knut and Alice Wallenberg, Family Erling Persson and Kempe Foundations, SciLifeLab, Stockholm University and Umeå University.

The molecular dynamics computations were partially enabled by resources provided by the National Academic Infrastructure for Supercomputing in Sweden (NAISS), i.e. the PDC Center for High Performance Computing at KTH Royal Institute of Technology, partially funded by the Swedish Research Council through grant agreement no. 2022-06725.

S.W. acknowledges funding from the Swedish Research Council (grant no. 2021-05101).

## Supporting information

Supporting file 1 containing Supplementary Fig.s1-11

## Data availability

The composite cryo-EM map used to reconstruct the PrPfr model has been deposited in EMDB (EMD-53916). The PrPfr model has been deposited in the Protein Data Bank (9RCC). Raw movies and final particle sets for the time points have been deposited in EMPIAR (EMPIAR-12874). Molecular dynamics simulations have been deposited in Zenodo (10.5281/zenodo.15796800).

## TOC graphic

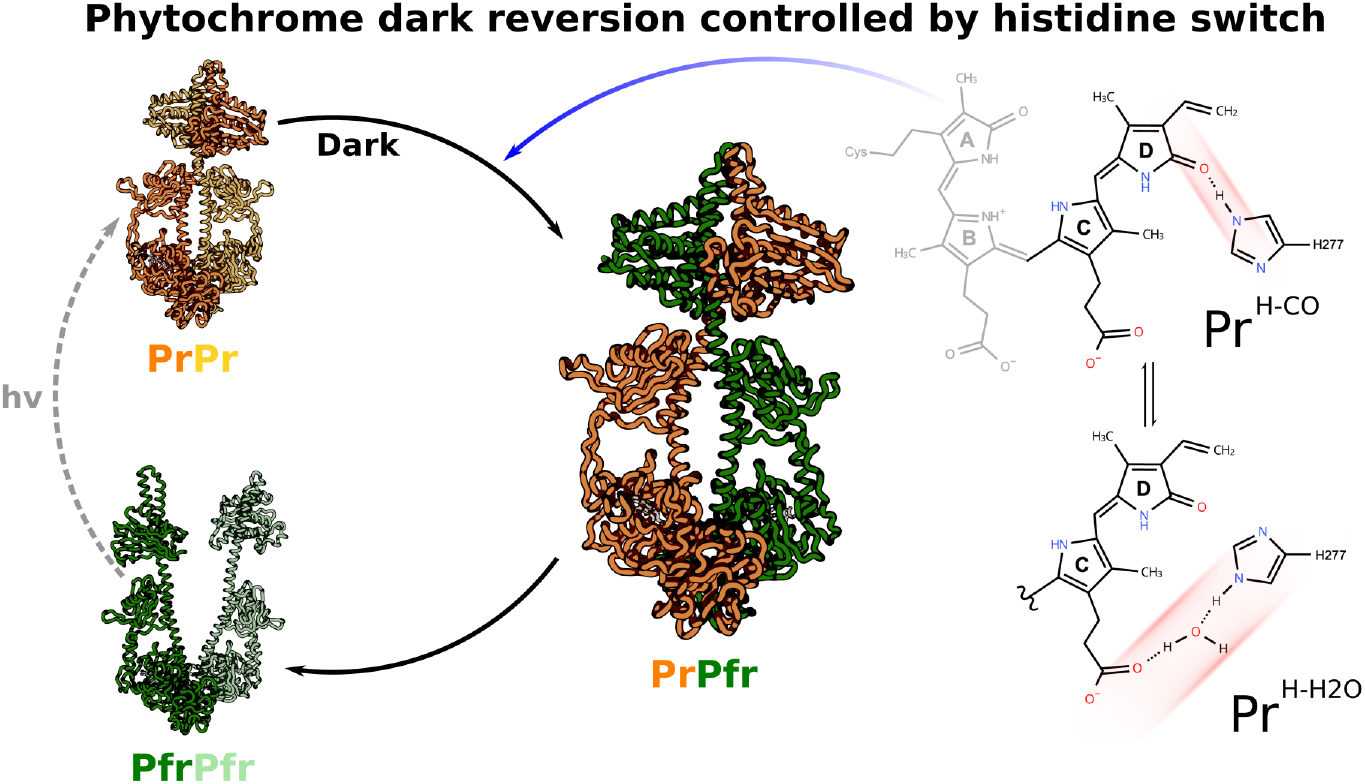

## Notes

### Competing Interest Statement

The authors have declared no competing interest.

### Summary of Updates

pH-dependent dark reversion rates were measured to support the claim that a histidine regulates the process (Fig. 3g); Results chapter was updated and the Discussion was revised to reflect the additional data.

## References

1. Butler, W. L., Norris, K. H., Siegelman, H. W. & Hendricks, S. B. DETECTION, ASSAY, AND PRELIMINARY PURIFICATION OF THE PIGMENT CONTROLLING PHOTORESPONSIVE DEVELOPMENT OF PLANTS. Proc Natl Acad Sci U S A 45, 1703–1708 (1959).

2. Hughes, J. et al. A prokaryotic phytochrome. Nature 386, 663 (1997).

3. Davis, S. J., Vener, A. V. & Vierstra, R. D. Bacteriophytochromes: phytochrome-like photoreceptors from nonphotosynthetic eubacteria. Science 286, 2517–2520 (1999).

4. Hughes, J. & Winkler, A. New Insight Into Phytochromes: Connecting Structure to Function. Annu Rev Plant Biol 75, 153–183 (2024).

5. Schäfer, E. & Schmidt, W. Temperature dependence of phytochrome dark reactions. Planta 116, 257–266 (1974).

6. Jung, J.-H. et al. Phytochromes function as thermosensors in Arabidopsis. Science 354, 886–889 (2016).

7. Casal, J. J. Photoreceptor signaling networks in plant responses to shade. Annu. Rev. Plant Biol. 64, 403–427 (2013).

8. Franklin, K. A. & Quail, P. H. Phytochrome functions in Arabidopsis development. J. Exp. Bot. 61, 11–24 (2010).

9. Legris, M., Ince, Y. Ç. & Fankhauser, C. Molecular mechanisms underlying phytochrome-controlled morphogenesis in plants. Nat. Commun. 10, 5219 (2019).

10. Hughes, J. Phytochrome cytoplasmic signaling. Annu Rev Plant Biol 64, 377–402 (2013).

11. Njimona, I., Yang, R. & Lamparter, T. Temperature effects on bacterial phytochrome. PLoS One 9, e109794 (2014).

12. Burgie, E. S. & Vierstra, R. Phytochromes: An Atomic Perspective on Photoactivation and Signaling. Plant Cell 26, 4568–4583 (2014).

13. Vierstra, R. D. & Davis, S. J. Bacteriophytochromes: new tools for understanding phytochrome signal transduction. Semin Cell Dev Biol 11, 511–521 (2000).

14. Takala, H., Edlund, P., Ihalainen, J. A. & Westenhoff, S. Tips and turns of bacteriophytochrome photoactivation. Photochem Photobiol Sci 19, 1488–1510 (2020).

15. Takala, H. et al. Signal amplification and transduction in phytochrome photosensors. Nature 509, 245–248 (2014).

16. Essen, L.-O., Mailliet, J. & Hughes, J. The structure of a complete phytochrome sensory module in the Pr ground state. Proc Natl Acad Sci U S A 105, 14709–14714 (2008).

17. Yang, X., Kuk, J. & Moffat, K. Crystal structure of Pseudomonas aeruginosa bacteriophytochrome: photoconversion and signal transduction. Proc Natl Acad Sci U S A 105, 14715–14720 (2008).

18. Gourinchas, G. et al. Long-range allosteric signaling in red light-regulated diguanylyl cyclases. Sci Adv 3, e1602498 (2017).

19. Björling, A. et al. Structural photoactivation of a full-length bacterial phytochrome. Science Advances (2016) doi:10.1126/sciadv.1600920.

20. Gourinchas, G., Heintz, U. & Winkler, A. Asymmetric activation mechanism of a homodimeric red light-regulated photoreceptor. Elife 7, (2018).

21. Li, H., Burgie, E. S., Gannam, Z. T. K., Li, H. & Vierstra, R. D. Plant phytochrome B is an asymmetric dimer with unique signalling potential. Nature 604, 127–133 (2022).

22. Burgie, E. S. et al. The structure of Arabidopsis phytochrome A reveals topological and functional diversification among the plant photoreceptor isoforms. Nat. Plants 9, 1116–1129 (2023).

23. Wahlgren, W. Y. et al. Structural mechanism of signal transduction in a phytochrome histidine kinase. Nat Commun 13, 7673 (2022).

24. Malla, T. N. et al. Photoreception and signaling in bacterial phytochrome revealed by single-particle cryo-EM. Sci Adv 10, eadq0653 (2024).

25. Bódizs, S., Mészáros, P., Grunewald, L., Takala, H. & Westenhoff, S. Cryo-EM structures of a bathy phytochrome histidine kinase reveal a unique light-dependent activation mechanism. Structure 32, 1952–1962.e3 (2024).

26. Wang, Z. et al. Light-induced remodeling of phytochrome B enables signal transduction by phytochrome-interacting factor. Cell 187, 6235–6250.e19 (2024).

27. Nagano, S. et al. Pr and Pfr structures of plant phytochrome A. Nature Communications 16, 1–12 (2025).

28. Björling, A. et al. Ubiquitous Structural Signaling in Bacterial Phytochromes. J Phys Chem Lett 6, 3379–3383 (2015).

29. Medzihradszky, M. et al. Phosphorylation of phytochrome B inhibits light-induced signaling via accelerated dark reversion in Arabidopsis. Plant Cell 25, 535–544 (2013).

30. Frankland, B. Biosynthesis and dark transformations of phytochrome. (1972) doi:10.5555/19730709563.

31. Anderson, G. R., Jenner, E. L. & Mumford, F. E. Temperature and pH studies on phytochrome in vitro. Biochemistry 8, 1182–1187 (1969).

32. Sweere, U. et al. Interaction of the response regulator ARR4 with phytochrome B in modulating red light signaling. Science 294, 1108–1111 (2001).

33. Eichenberg, K. et al. Arabidopsis phytochromes C and E have different spectral characteristics from those of phytochromes A and B. FEBS Lett 470, 107–112 (2000).

34. Yi, C. et al. Plant Phytochrome Interactions Decode Light and Temperature Signals. Plant Cell 36, 4819–4839 (2024).

35. Legris, M. et al. Phytochrome B integrates light and temperature signals in Arabidopsis. Science 354, 897–900 (2016).

36. Hennig, L. & Schäfer, E. Both subunits of the dimeric plant photoreceptor phytochrome require chromophore for stability of the far-red light-absorbing form. J Biol Chem 276, 7913–7918 (2001).

37. Klose, C. et al. Systematic analysis of how phytochrome B dimerization determines its specificity. Nat Plants 1, 15090 (2015).

38. Prodhan, S. et al. Detection of a hybrid PrPfr state in the dark reversion of a bathy phytochrome indicates inter-dimer allostery. Phys. Chem. Chem. Phys. 27, 20279–20287 (2025).

39. Altoè, P. et al. Deciphering intrinsic deactivation/isomerization routes in a phytochrome chromophore model. J. Phys. Chem. B 113, 15067–15073 (2009).

40. Lagarias, J. C. & Rapoport, H. Chromopeptides from phytochrome. The structure and linkage of the PR form of the phytochrome chromophore. J. Am. Chem. Soc. 102, 4821–4828 (1980).

41. Villegas-Escobar, N. & Matute, R. A. The Keto-Enol Tautomerism of Biliverdin in Bacteriophytochrome: Could it Explain the Bathochromic Shift in the Pfr Form? Photochem Photobiol 97, 99–109 (2021).

42. Velazquez Escobar, F. et al. A protonation-coupled feedback mechanism controls the signalling process in bathy phytochromes. Nat Chem 7, 423–430 (2015).

43. Escobar, F. J. V. Vibrational Spectroscopy of Phytochromes and Phytochrome-Related Photoreceptors. (2015).

44. Stiller, J. B. et al. Structure determination of high-energy states in a dynamic protein ensemble. Nature 603, 528–535 (2022).

45. Song, C. et al. Two ground state isoforms and a chromophore D-ring photoflip triggering extensive intramolecular changes in a canonical phytochrome. Proc Natl Acad Sci U S A 108, 3842–3847 (2011).

46. Kim, P. W., Rockwell, N. C., Martin, S. S., Clark Lagarias, J. & Larsen, D. S. Heterogeneous Photodynamics of the Pfr State in the Cyanobacterial Phytochrome Cph1. (2014) doi:10.1021/bi5005359.

47. Yang, Y. et al. Active and silent chromophore isoforms for phytochrome Pr photoisomerization: An alternative evolutionary strategy to optimize photoreaction quantum yields. Struct Dyn 1, 014701 (2014).

48. Lim, S. et al. Correlating structural and photochemical heterogeneity in cyanobacteriochrome NpR6012g4. Proc Natl Acad Sci U S A 115, 4387–4392 (2018).

49. Mukherjee, S., Jemielita, M., Stergioula, V., Tikhonov, M. & Bassler, B. L. Photosensing and quorum sensing are integrated to control Pseudomonas aeruginosa collective behaviors. PLoS Biol 17, e3000579 (2019).

50. Multamäki, E. et al. Comparative analysis of two paradigm bacteriophytochromes reveals opposite functionalities in two-component signaling. Nat Commun 12, 4394 (2021).

51. Huber, C., Strack, M., Diller, R. & Frankenberg-Dinkel, N. Analyzing In Vitro Autokinase Activity of Microbial Phytochromes Under Well-Defined Light Conditions. Methods Mol Biol 2970, 285–295 (2026).

52. Takala, H., Björling, A., Linna, M., Westenhoff, S. & Ihalainen, J. A. Light-induced Changes in the Dimerization Interface of Bacteriophytochromes. J Biol Chem 290, 16383–16392 (2015).

53. Isaksson, L. et al. Signaling Mechanism of Phytochromes in Solution. Structure 29, 151–160.e3 (2021).

54. Yang, Y. et al. Ultrafast proton-coupled isomerization in the phototransformation of phytochrome. Nat Chem 14, 823–830 (2022).

55. Akke, M. & Weininger, U. NMR Studies of Aromatic Ring Flips to Probe Conformational Fluctuations in Proteins. J Phys Chem B 127, 591–599 (2023).

56. Chenchiliyan, M. et al. Ground-state heterogeneity and vibrational energy redistribution in bacterial phytochrome observed with femtosecond 2D IR spectroscopy. J Chem Phys 158, 085103 (2023).

57. Salvadori, G. & Mennucci, B. Analogies and Differences in the Photoactivation Mechanism of Bathy and Canonical Bacteriophytochromes Revealed by Multiscale Modeling. J Phys Chem Lett 15, 8078–8084 (2024).

58. Mathes, T. et al. Femto- to Microsecond Photodynamics of an Unusual Bacteriophytochrome. J Phys Chem Lett 6, 239–243 (2015).

59. Klose, C., Nagy, F. & Schäfer, E. Thermal reversion of plant phytochromes. Mol. Plant 13, 386–397 (2020).

60. Croll, T. I. ISOLDE: a physically realistic environment for model building into low-resolution electron-density maps. Acta Crystallogr D Struct Biol 74, 519–530 (2018).

61. Anandakrishnan, R., Aguilar, B. & Onufriev, A. V. H++ 3.0: automating pK prediction and the preparation of biomolecular structures for atomistic molecular modeling and simulations. Nucleic Acids Res 40, W537–41 (2012).

62. Gordon, J. C. et al. H++: a server for estimating pKas and adding missing hydrogens to macromolecules. Nucleic Acids Res 33, W368–71 (2005).

63. Myers, J., Grothaus, G., Narayanan, S. & Onufriev, A. A simple clustering algorithm can be accurate enough for use in calculations of pKs in macromolecules. Proteins 63, 928–938 (2006).

64. H++ (web-based computational prediction of protonation states and pK of ionizable groups in macromolecules). http://newbiophysics.cs.vt.edu/H++/.

65. Maier, J. A. et al. ff14SB: Improving the Accuracy of Protein Side Chain and Backbone Parameters from ff99SB. J Chem Theory Comput 11, 3696–3713 (2015).

66. Jorgensen, W. L., Chandrasekhar, J., Madura, J. D., Impey, R. W. & Klein, M. L. Comparison of simple potential functions for simulating liquid water. J. Chem. Phys. 79, 926–935 (1983).

67. El Hage, K., Hédin, F., Gupta, P. K., Meuwly, M. & Karplus, M. Valid molecular dynamics simulations of human hemoglobin require a surprisingly large box size. Elife 7, (2018).

68. Gapsys, V. & de Groot, B. L. Comment on ‘Valid molecular dynamics simulations of human hemoglobin require a surprisingly large box size’. Elife 8, (2019).

69. Case, D. A. et al. Amber Tools. Journal of Chemical Information and Modeling (2023) doi:10.1021/acs.jcim.3c01153.

70. Hub, J. S., de Groot, B. L., Grubmüller, H. & Groenhof, G. Quantifying Artifacts in Ewald Simulations of Inhomogeneous Systems with a Net Charge. J Chem Theory Comput 10, 381–390 (2014).

71. Wallnoefer, H. G., Handschuh, S., Liedl, K. R. & Fox, T. Stabilizing of a globular protein by a highly complex water network: a molecular dynamics simulation study on factor Xa. J Phys Chem B 114, 7405–7412 (2010).

72. Salomon-Ferrer, R., Götz, A. W., Poole, D., Le Grand, S. & Walker, R. C. Routine Microsecond Molecular Dynamics Simulations with AMBER on GPUs. 2. Explicit Solvent Particle Mesh Ewald. J Chem Theory Comput 9, 3878–3888 (2013).

73. Miyamoto, S. & Kollman, P. A. Settle: An analytical version of the SHAKE and RATTLE algorithm for rigid water models. J. Comput. Chem. 13, 952–962 (1992).

74. Adelman, S. A. & Doll, J. D. Generalized Langevin equation approach for atom/solid-surface scattering: General formulation for classical scattering off harmonic solids. J. Chem. Phys. 64, 2375–2388 (1976).

75. Åqvist, Johan, et al. “Molecular dynamics simulations of water and biomolecules with a Monte Carlo constant pressure algorithm.” Chemical physics letters 384.4-6 (2004): 288–294.

76. Roe, D. R. & Cheatham, T. E., 3rd. PTRAJ and CPPTRAJ: Software for Processing and Analysis of Molecular Dynamics Trajectory Data. J Chem Theory Comput 9, 3084–3095 (2013).

77. VMD: Visual molecular dynamics. Journal of Molecular Graphics 14, 33–38 (1996).

78. Fernández-Quintero, M. L. et al. The influence of antibody humanization on shark variable domain (VNAR) binding site ensembles. Front Immunol 13, 953917 (2022).

