## Supporting Information for "Chemical mechanism of allosteric and asymmetric dark reversion in a bacterial phytochrome uncovered by cryo-EM"


### Authors

Szabolcs Bódizs[1^], Anna-Lena M. Fischer[1^], Miklós Cervenak[1], Sayan Prodhan[1], Michal Maj[2], and Sebastian Westenhoff[1,*]

### Affiliations

[1] Department of Chemistry - BMC, Biochemistry, Uppsala University, 75123 Uppsala, Sweden

[2]Department of Chemistry - Ångström, Uppsala University, 75123 Uppsala, Sweden

[^]Equal contribution

[*]Lead contact

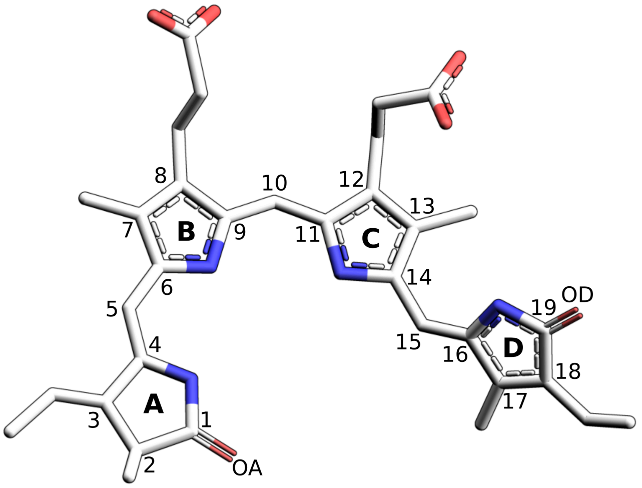


***Figure S1. The structure and atomic numbering of the biliverdin chromophore.*** *"OD" labels the oxygen on the D ring, proposed to undergo keto-enol tautomerization.*

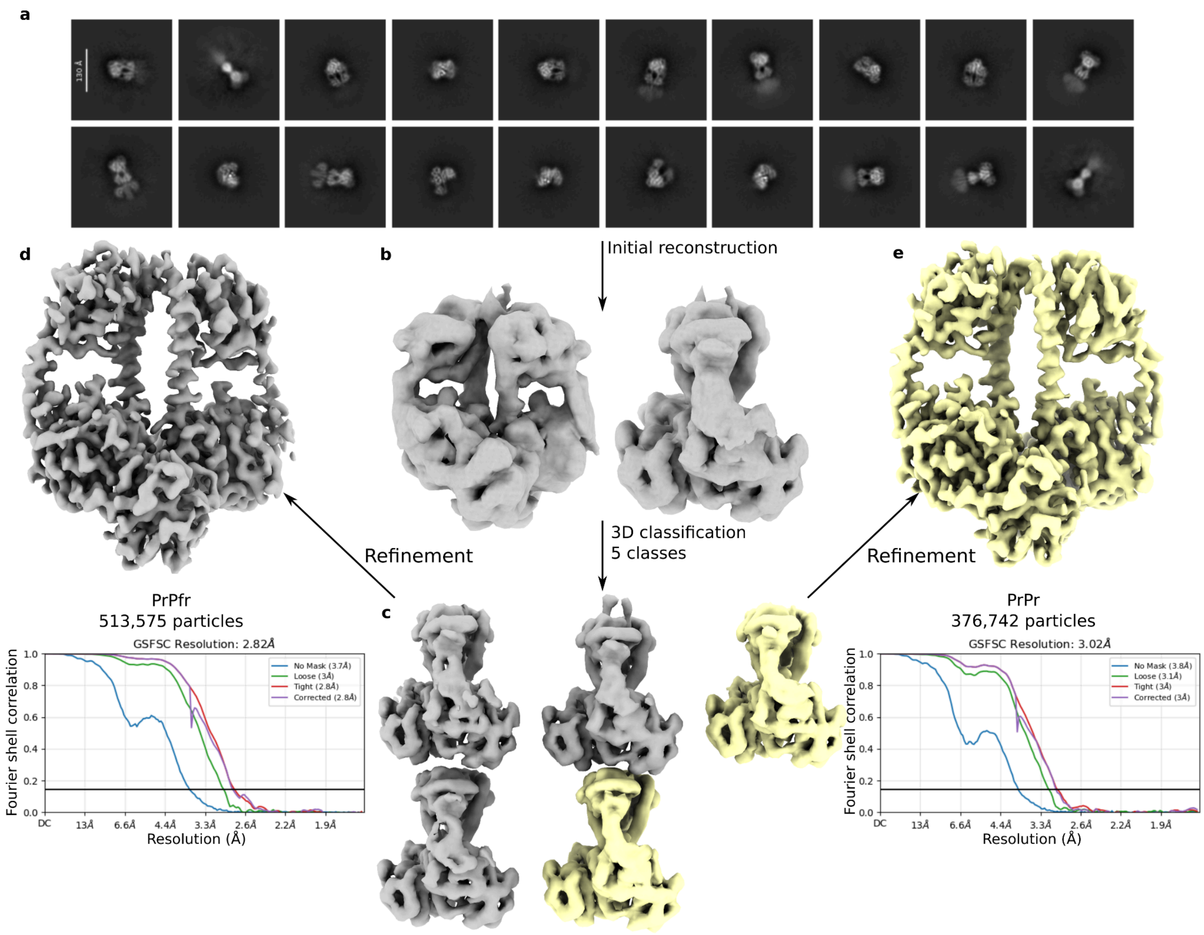


***Figure S2. Reconstruction of the PrPr and PrPfr states from cryo-EM data collected 120 s into the dark reversion process.*** *2D classes show a closed dimer majority, with less than 0.1% of the particles forming open dimer classes (a). Consensus refinement (b) showed an intermediate position for tongue*$_B$*, which was separated into PrPr (yellow) and PrPfr (grey) classes using 3D classification (c). The two states were then locally refined using the photosensory module as a mask (d, e).*

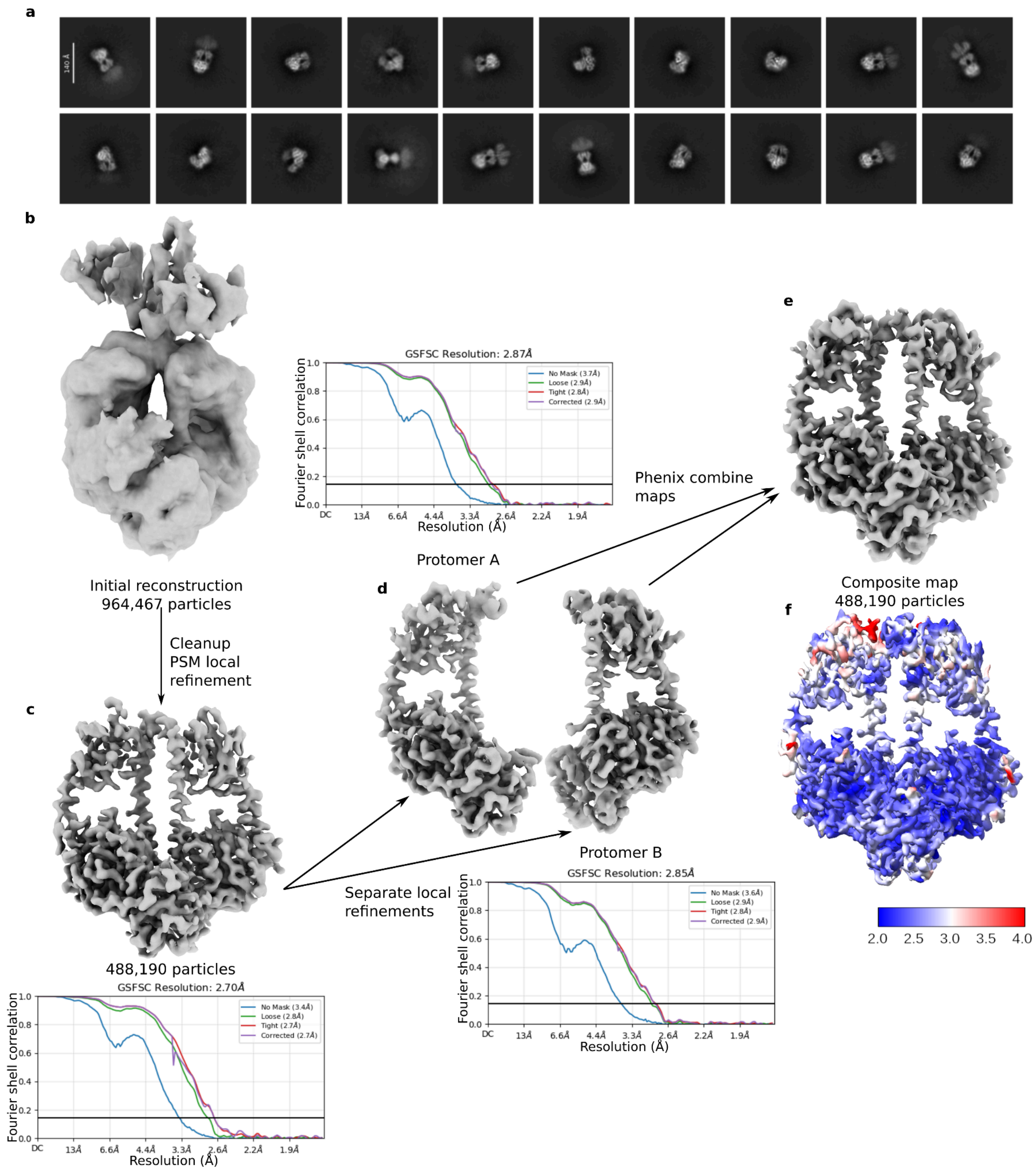


***Figure S3. Reconstruction of the PrPfr state from cryo-EM data collected 300 s into the dark reversion process.*** *2D classes show a closed dimer majority (the PrPr and PrPfr states are indistinguishable at this level of detail), with less than 2% of the particles forming open dimer classes (a). Consensus refinement (b) reveals the orientation of the output module at low resolution, but is lost when locally refining the PSM (c). Separate local refinements of the two protomers (d) improve the resolution in the binding pocket (which was blurred due to minor flexibility of the dimer interface). The map used for model building is the composite (e) of the two local refinements, with local resolution shown in (f).*

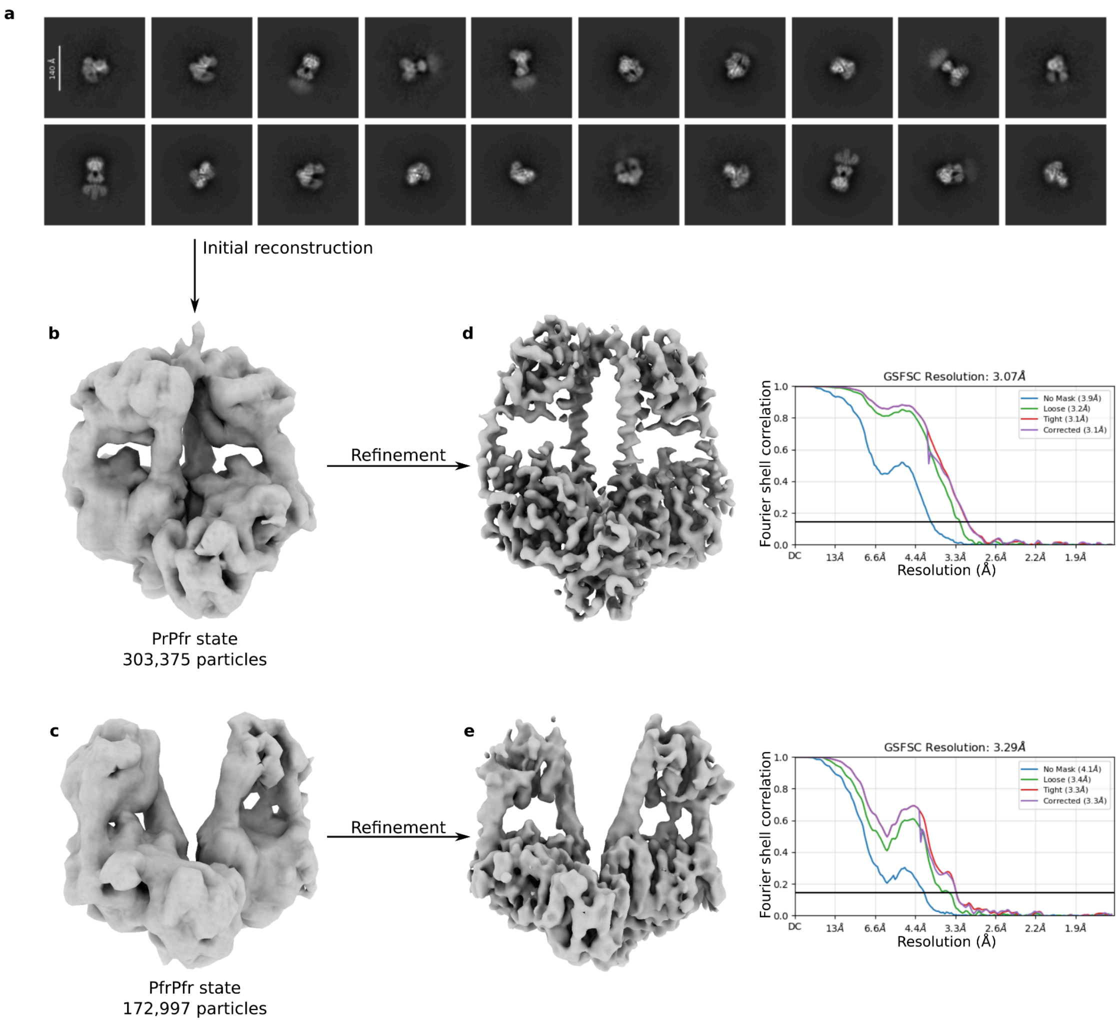


***Figure S4. Reconstruction of the PrPfr and PfrPfr states from cryo-EM data collected 960s into the dark reversion process.*** *2D classification (a) identifies a mixture of open and closed states. A PrPfr (b) and a PfrPfr (c) state were reconstructed and subsequently refined (d, e). The resolution of the PfrPfr map is lower due to a lower particle number and increased flexibility, but the conformation of the PHY tongue confirms its photochromic state.*

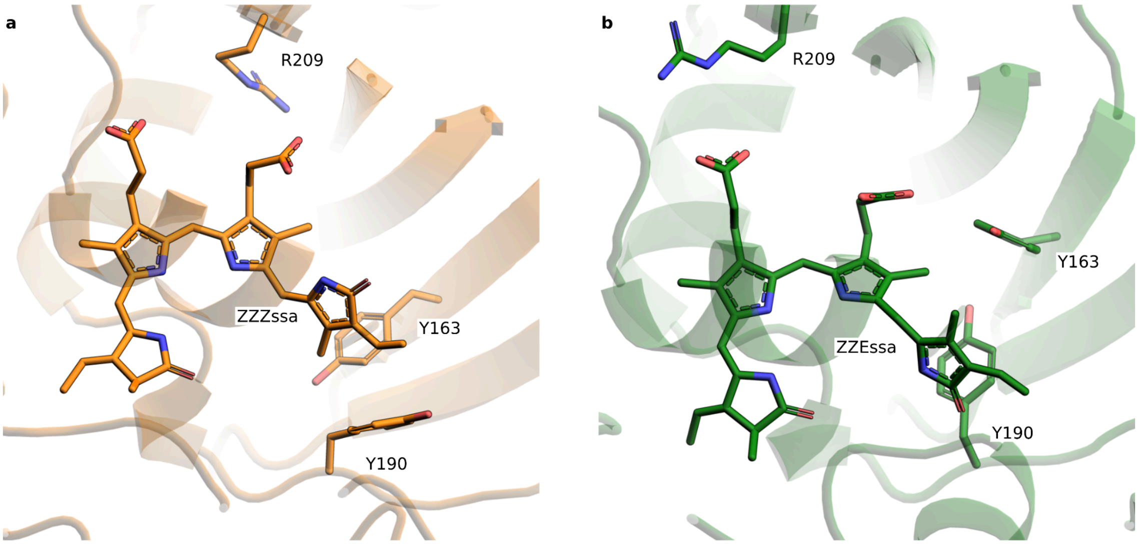


***Figure S5. The chromophore binding pocket of the PrPfr hybrid state.*** *Pr in protomer A and Pfr in protomer B are shown in (a) and (b), respectively. The chromophore isomerizes from ZZZ (Pr) to ZZE (Pfr). The residues Y163, Y190 and R209 rotate, rearranging the binding pocket.*

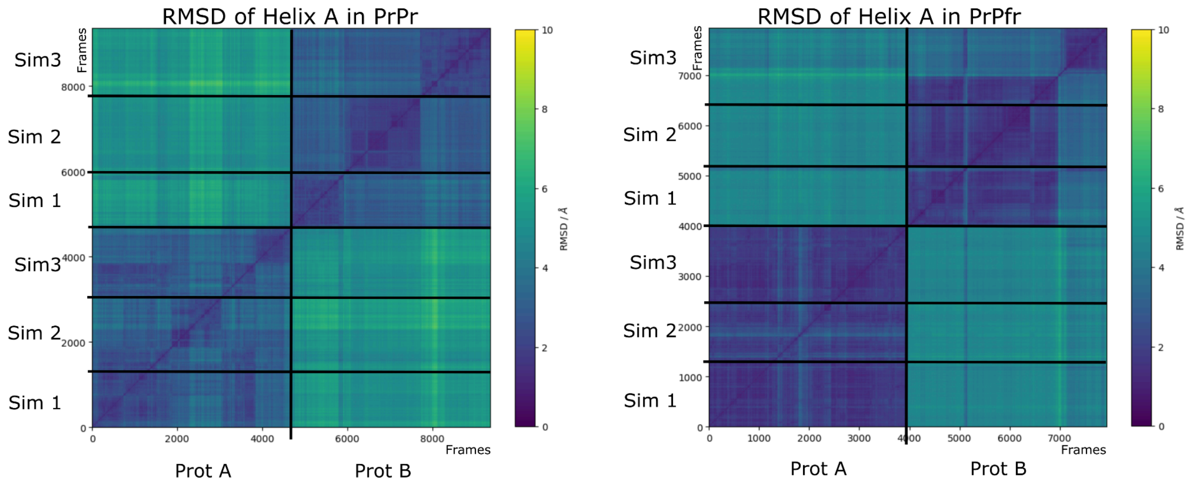


***Figure S6. 2D-RMSD plots of the simulations for PrPr (left) and PrPfr (right)**, to show that the distortion in helix A is only present in protomer A and not in protomer B. The simulations were aligned on the spine and helix B (residue 286-304 and 139-155) to pinpoint the helix A (residue 123-136) conformational changes in the three-helix bundle by RMSD on the C-alpha atoms. A higher RMSD in the region where protomer A gets compared to protomer B while in between the protomers the RMSD is rather low shows that the helix adopts the same conformations throughout the simulations of PrPr and also in PrA of PrPfr. Only in the already reverted protomer we find higher flexibility and different states which Helix a can adopt (different from the two identified).*

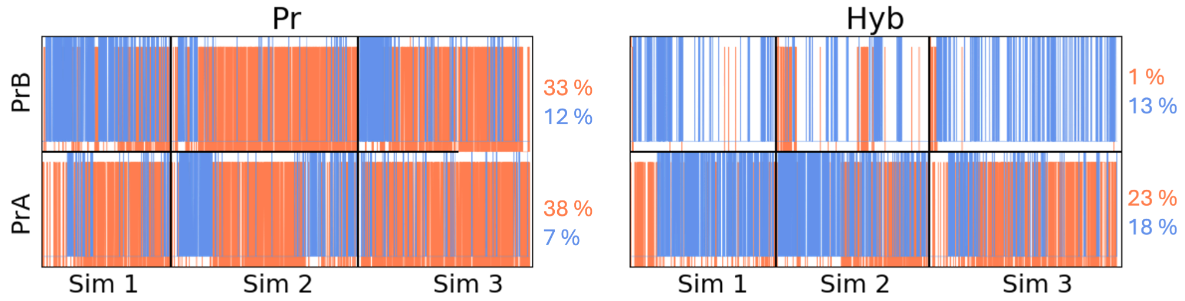


***Figure S7. Time evolution of the hydrogen bond formation for all simulations for each protomer respectively.*** *The color scheme is set according to Fig. 3e: His277-OD in red and His-H2O in blue. This shows convergence for this interaction network since we see many transitions over the course of all simulations.*

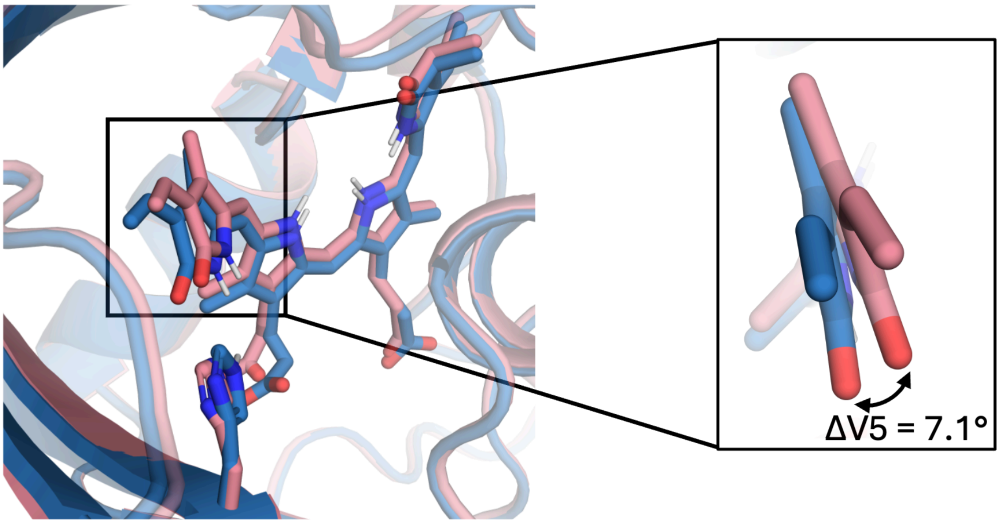


***Fig. S8. The conformation of the D-ring is controlled by the hydrogen bonding of His277.*** *Overlay of the average structures (derived from the MD simulations) of the "H-OD" (red) and "H-H2O" (blue) arrangements. We observe a notable shift in the V5 torsion (NC-C14-C15-C16, see numbering in S1) from -145.5 ("H-OD") to -152.6 ("H-H2O").*

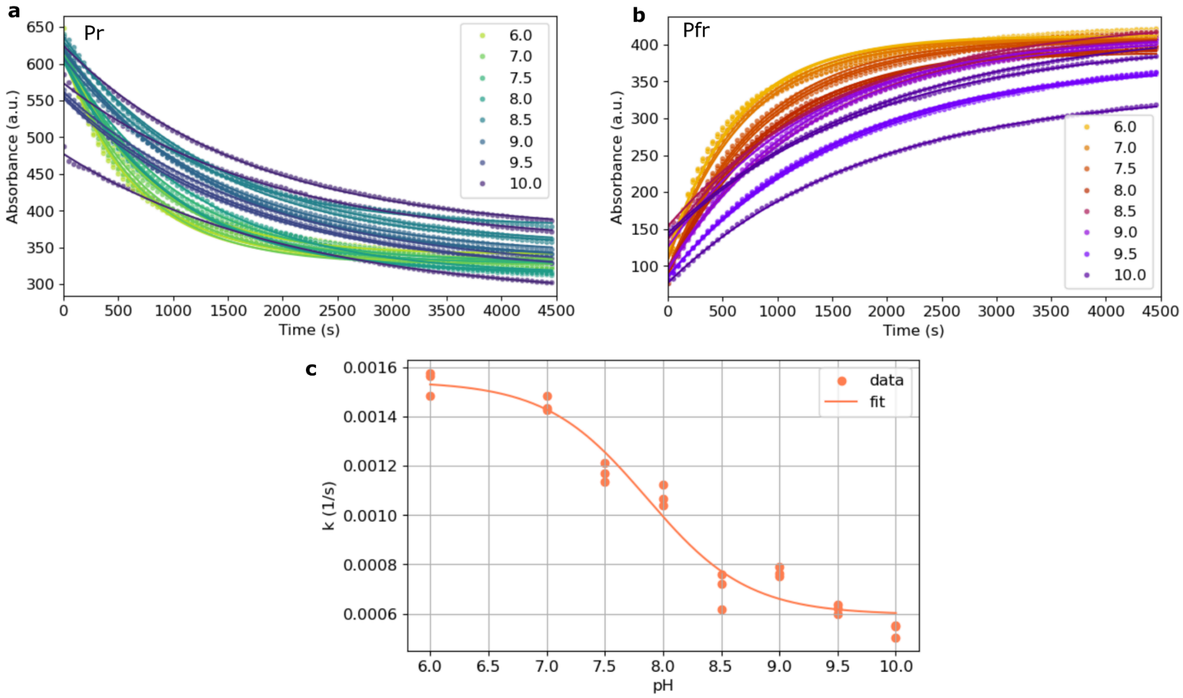


***Figure S9. pH dependence of dark reversion according to Pr decay and Pfr growth.*** *(a) and (b) show the absorbance decay of Pr and growth of Pfr at different pH values between 6 and 10, three repeats each with the respective exponential fits. In (c) is the sigmoidal fit with a Henderson-Hasselbach-like function (see Methods) of the Pfr growth rate constant, resulting in a pKa of 7.9.*

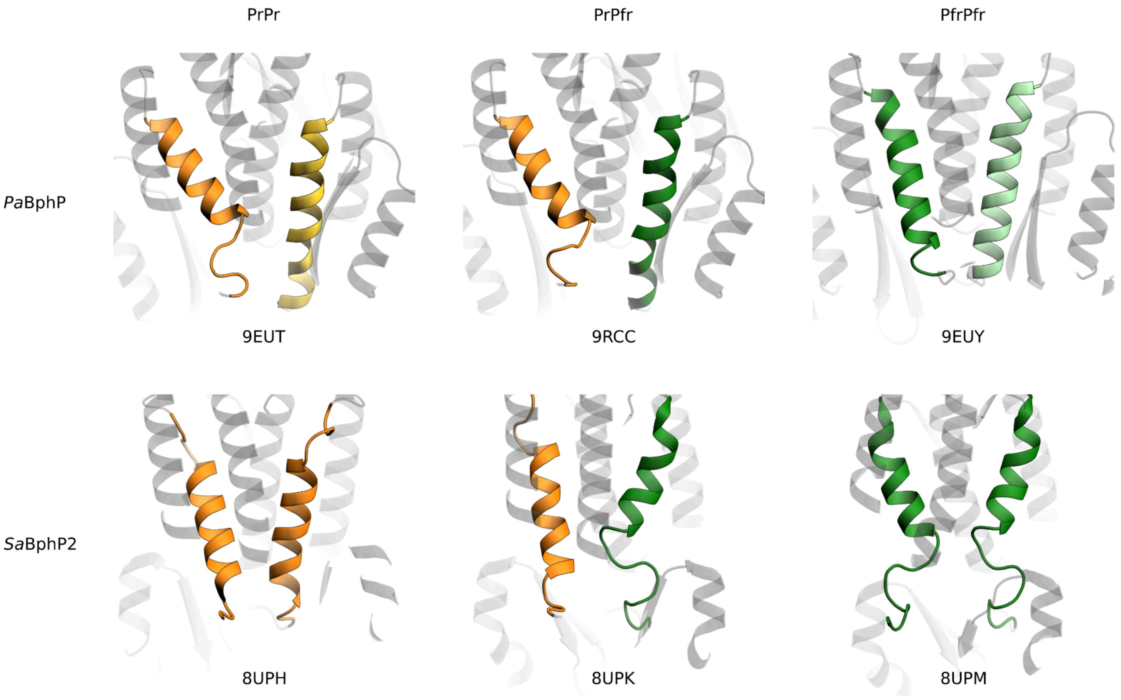


***Figure S10. The states of GAF helix A in* Pa*BphP and* Sa*BphP2 in their PrPr, PrPfr and PfrPfr forms.*[1,2]** Pa*BphP is asymmetric in PrPr, which carries over to the PrPfr state. The asymmetry in PrPfr in* Sa*BphP2 is simply a product of the different photochromic states and does not seem to be present in either ground state. Helix A (highlighted) is defined as residues 118-136 in* Pa*BphP, and residues 114-134 in* Sa*BphP2.*

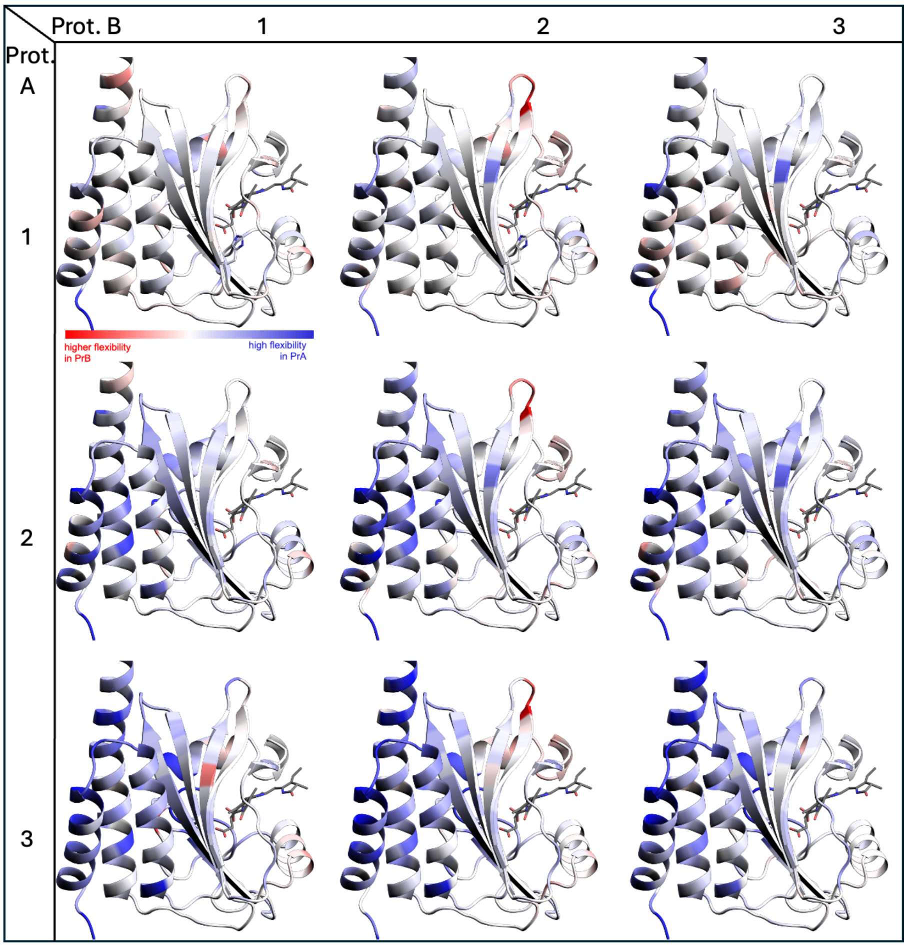


***Figure S11. Consistency in dynamics difference of the two protomers.*** *RMSF difference of each protomer A to each protomer B of all three PrPr simulations. While protomer A of simulation 1 only shows small differences tending to a higher flexibility, we clearly see this effect for all other simulations.*

### Supporting references


1. Bódizs, S., Mészáros, P., Grunewald, L., Takala, H. & Westenhoff, S. Cryo-EM structures of a bathy phytochrome histidine kinase reveal a unique light-dependent activation mechanism. *Structure* **32**, 1952–1962.e3 (2024).
2. Malla, T. N. *et al.* Photoreception and signaling in bacterial phytochrome revealed by single-particle cryo-EM. *Sci Adv* **10**, eadq0653 (2024).
